# Sex-specific functions of duetting in the Galápagos Yellow Warbler (Setophaga petechia aureola)

**DOI:** 10.64898/2026.07.10.737681

**Authors:** Alper Yelimlieş, Çağlar Akçay, Sonia Kleindorfer

## Abstract

In over a thousand avian species, pairs coordinate vocalizations to form duets. Songbirds may use calls (simple, non-learned vocalizations) and songs (complex, learned vocalizations) to answer their partners. In Galápagos Yellow Warblers (*Setophaga petechia aureola*), song answers in duets appear to have evolved recently in addition to older call answers. Here, we explored the structure and functions of duets in an observational study to understand why song answers in duets may have evolved. We first described the temporal structure of duets initiated by either sex. We then tested whether the distance between pair members, countersinging with neighbours, and timing relative to aggressive encounters predicted answering probability for each sex. Finally, we tested whether the behavioral context of call and song answers differed. We found that most duets were initiated by males, and call answers were almost exclusively given by females. Female answers were also more tightly coordinated than those of males. Female answering probability decreased with increasing distance between pair members, whereas male answering probability did not. In addition, females were less likely to answer with songs when farther from their mate. Females’ answering probability also increased slightly when their mate’s initiation was preceded by a neighbor’s song. Our results show some evidence in support of the joint resource defence hypothesis for female answers. On the other hand, the significance of song in female answers may be related to some other functions that we did not test here, such as signaling quality or commitment to the pair bond.

**Lay summary:** Who turns a song into a duet? In Galápagos Yellow Warblers, it is usually the female: she answers her mate’s song with a tightly timed call or song. Male replies were rarer and less coordinated. While male replies didn’t seem to have a function, female replies were used in territorial disputes with neighbours and may indicate joint ownership of a territory. Female song replies, on the other hand, may signal commitment to the pair bond.

---

Duets, temporally coordinated acoustic signals produced by two individuals, have been documented across a wide range of animal taxa (Thorpe et al. 1972; Tauber and Pener 2000; Clink et al. 2020; Legett et al. 2021). In birds, more than 1600 species spanning a broad taxonomic range produce duets (Tobias et al. 2016). Although their antiphonal quality and temporal precision inspired early research on duetting (Thorpe, 1963), duet structure varies considerably both among and within species, including overlapping or loosely coordinated phrases (Hall 2009; Dahlin and Benedict 2014).

The structure of the duets is often linked to their function. For instance, louder duets and duets containing sex-specific phrases tend to be externally directed, reflecting conflict or cooperation between the pair (Dahlin and Benedict 2014). While precise and alternating phrases may function in cooperative territory defense, overlapping phrases may mask the mate’s vocalization as a form of mate guarding (Tobias and Seddon 2009). Because the costs and benefits of duetting may differ between pair members, individual-level behaviors should be examined separately to understand emergent pair-level properties of duets (Hall 2009; Logue and Krupp 2016).

Paired mates may coordinate their acoustic signals for multiple non-mutually exclusive functions to communicate with extrapair receivers and each other, such as joint territorial defense and maintaining contact (reviewed in detail by Hall, 2004). According to the joint resource defense hypothesis, duets are cooperative signals directed to extrapair receivers that are used to advertise territories to keep out conspecifics or mediate agonistic interactions with intruders.

Numerous studies provide strong support for this hypothesis across duetting species (Logue 2005; Dahlin and Benedict 2014). In contrast, the contact maintenance hypothesis proposes that duets function primarily in within-pair communication, allowing mates to maintain contact by initiating and answering duets and thereby localize one another, especially in densely vegetated habitats (Thorpe, 1963).

Although experimental approaches have revealed much about the adaptive significance of duetting (Levin 1996a; Levin 1996b), observational studies are essential for understanding how duets function under natural conditions. Such studies have been pivotal to advancing our understanding that duets occur in different contexts, may serve multiple functions, and these functions can be sex specific. For example, in rufous-and-white wrens (*Thryothorus rufalbus*), pair localization using a recording array revealed that individuals use their mates’ answers to localize and approach them (Mennill and Vehrencamp 2008). Similarly, radio-tracking of the black-bellied wrens (*Pheugopedius fasciatoventris*) showed that answering increased the initiator’s likelihood of approaching and that duet answering was more likely when the pair was already close to each other (Logue 2007). Moreover, multiseasonal observations of Venezuelan troupials (*Icterus icterus*) indicated that, although both sexes answer to their mates to defend territories year-round, males may also use duets for mate guarding during the breeding season (Odom et al. 2017). Together, these findings demonstrate that observing the natural context of duetting can provide important insights into duet function for each sex.

As duets are not limited to songbirds, contributions by pair members are not exclusively learned vocalizations (i.e., songs); they may also include calls and non-vocal sonations (Farabaugh 1982). In some songbird families, both sexes commonly sing; however, parulids, where duetting and female song appear to have evolved more recently (Mitchell et al. 2019, Najar and Benedict 2015), offer an interesting case study for examining how calls and songs are used in duets. In some species of the genus *Myiothlypis,* in which female song is frequent, duets consist of songs from both sexes, like in the Choco warbler (*Myiothlypis chlorophry*s; Curso et al., 2022). In the genus *Setophaga,* however, with infrequent or absent female song, duets can comprise male song initiations answered by female calls (Spector 1991; Krause et al. 2026). We recently described Galápagos Yellow Warbler (*Setophaga petechia aureola*) duets in which both males and females combine songs (see Figure 1; Yelimlieş et al., 2026). Moreover, females of both Galápagos Yellow Warblers and of the sister species Northern Yellow Warblers (*Setophaga aestiva*) utter *chip* calls at the end of their mate’s song to form duets (Snow 1966; Spector 1991). Although Northern Yellow Warbler females have been reported to sing rare solo songs, they don’t combine them with their partners’ songs to sing duets (Hobson and Sealy 1990). To our knowledge, the Galápagos Yellow Warbler is therefore the only species in the genus *Setophaga* in which frequent song duets have been documented. This raises the question: what selective pressures favoured the evolution of song answering in duets?

**Figure 1.**
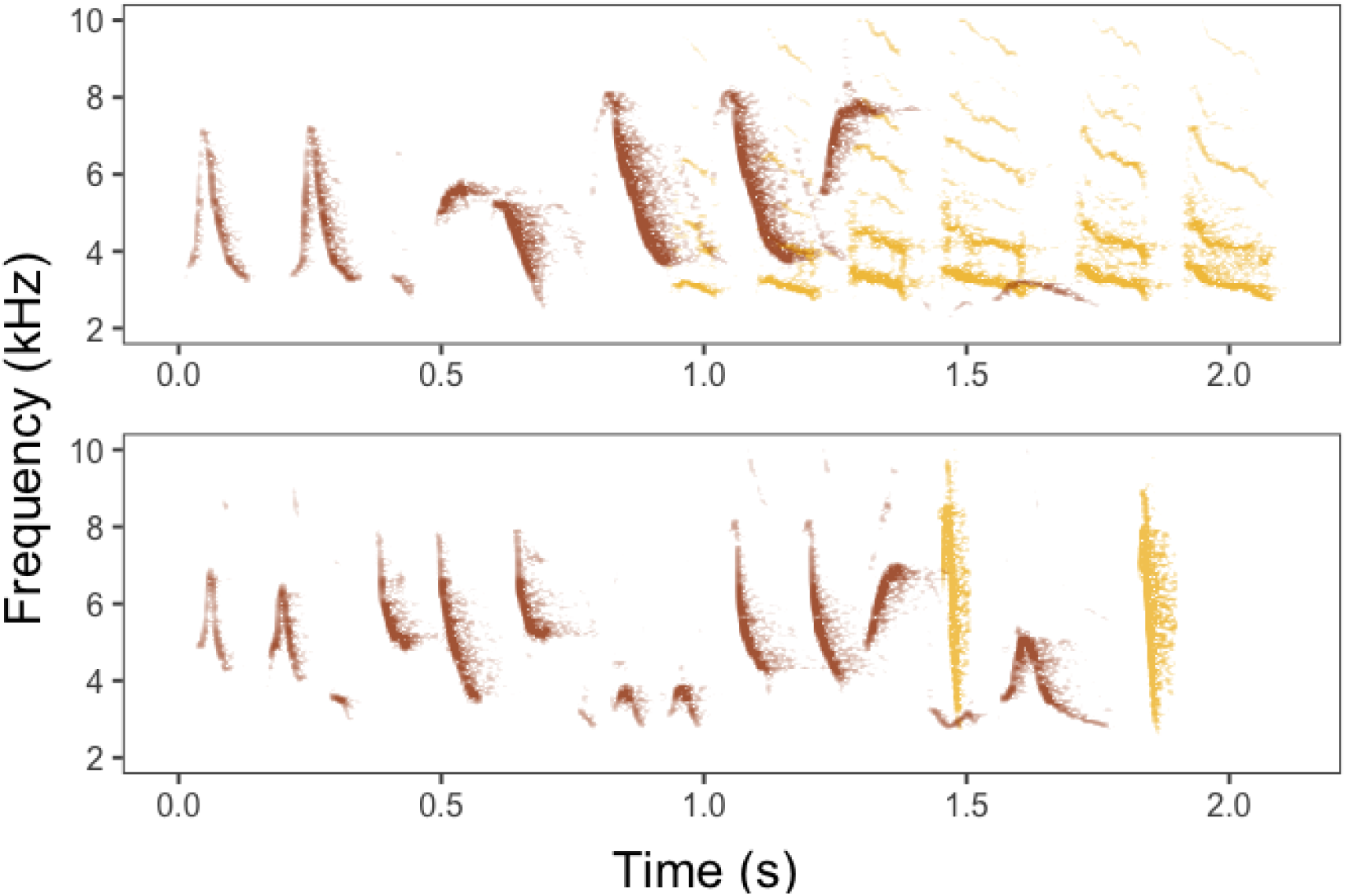
Spectrograms of the two duet types of Galápagos Yellow Warblers. In these examples, both duets are initiated by a male song for simplicity. In the top spectrogram, the female answers with a song, whereas in the bottom spectrogram, she answers with two *chip* calls. Male songs are presented in brown, female vocalizations are in yellow. For illustration purposes, duets are created by combining solo vocalizations of each sex with the natural temporal latency of answers. Spectrograms are created using the ggspectro function (window length = 1024, window type = hann, overlap = 90) from the seewave package in R (Sueur et al. 2008).

In the present study, we aimed to uncover the potential functions of duetting and different answer types in each sex by observing the natural contexts in which duets occurred. We thus recorded the vocalizations of pairs, aggressive interactions, countersinging with neighbors, and distances between pair members. We first describe the temporal structure of duets and then tested the functions of duet answering for each sex, with particular emphasis on song versus call answers. We predicted that 1) If answering functions in mediating aggressive encounters, answering probability should peak shortly before or during aggressive encounters. Alternatively, 2) if answering functions as a post-contest signal, answering probability should peak shortly after aggressive encounters. 3) If answering functions in territory boundary disputes, answers should be more likely when countersinging with neighbors. 4) If answering functions in contact maintenance between the pair members, answers should be more likely when pair members are further apart. Because the costs and benefits of answering may differ between the sexes, we analyzed males and females separately. Finally, if song answering evolved specifically for one or more of these contexts, we expected the probability of answering with a song to be higher than the probability of answering with a *chip* call in those contexts.

## Methods

### Study species

The Galápagos Yellow Warbler is a subspecies of the Mangrove Yellow Warbler endemic to the Galápagos and Cocos Islands. In contrast to Northern Yellow Warblers, Mangrove Yellow Warblers do not migrate and are territorial year-round. Unlike their mainland counterparts inhabiting Central and South America, Galápagos Yellow Warblers are not mangrove specialists and are abundant in both highland forests and lowland shrubs of the Galápagos Archipelago. Pair-bonded males and females maintain year-round territories and often remain together for multiple years (Snow 1966; Yelimlieş et al. 2026). Galápagos Yellow Warblers are sexually dimorphic in both plumage and song. Males have a rufous crown and prominent chest streaking, whereas females lack the crown and have little or no streaking. Female songs also sound harsher and more repetitive than male songs to the human ear.

During simulated territorial intrusions, both sexes defend the territory; however, females don’t participate in defense when they have an active nest (Yelimlieş et al. 2026). Females rarely respond to their mate’s song with a song of their own during the breeding season, but female song answers are common in the non-breeding season. Females also respond to their mate’s song with “chip” calls in both breeding and non-breeding seasons. Territorial behavior of the Galápagos subspecies is poorly described outside experimental contexts, unlike that of the Northern sister species (Ficken & Ficken, 1965; Valdez-Juárez et al., 2025). In the Costa Rican Mangrove Yellow Warbler (*S. p. xanthotera*), both sexes repel conspecific intruders, whereas females primarily chase heterospecifics (Barrantes 1998). We similarly observed both sexes chasing intruders in the Galápagos, with these brief interactions accompanied by a rapid series of “chip” and “dzeet” calls. The latter is a short frequency-modulated call similar to that described in Northern Yellow Warblers.

### Data collection and processing

We collected data from ten pairs between 27 June and 8 July 2024 on Floreana Island, where the population has been a part of a long-term study of landbirds (Kleindorfer et al. 2021; Hohl et al. 2025). We selected pairs with known territory boundaries by observing their singing posts and ensured that both members of the pair were color-banded before observations. All territories were in Puerto Velazco Ibarra, the island’s only settlement. The habitat consisted mainly of low shrubs, allowing easier visual tracking of individuals than in the dense *Scalesia* forests of the highlands. We observed the pairs in one-hour blocks for a total of 17 hours; all pairs except three were observed for two blocks on different days. Each observation was carried out by two researchers, with one observer following the male and the other following the female to minimize behavioral omissions. Observers continuously narrated the behavior of their focal individual, including vocalizations, agonistic encounters, and any singing neighbors. They also independently estimated the distance between pair members and agreed on a final estimate in real time. Narrations and vocalizations were recorded using a Zoom F3 recorder connected to a Sennheiser MKE 600 or ME 66/K6 shotgun microphone.

We then used Audacity version 3.7.1 to annotate our recordings of the observations. We first temporally aligned both recordings from an observation session. From the recordings, we extracted three types of vocal events: solo songs (i.e., an unanswered duet initiation), song answers, and call answers. Following our previous approach, we defined a duet as a song by one pair member answered by another song, or a call bout by the other pair member, with the answer starting after the initiation and not exceeding the end of the answer by more than two seconds. Call bouts were composed of calls with inter-onset intervals of less than 1 second. For each of these vocal events, we coded whether a neighbor song was present within the 10-second window preceding the event onset. This window allows for song exchanges between neighbors as it approximates the rate at which Northern Yellow Warblers sing (Spector 1991), and consistent with our observations of the gap between two songs of a male Galápagos Yellow Warblers. We also coded the distance between pair members at the time of the event and the relative time difference between the event onset and the closest aggressive encounter. These encounters were brief, and were accompanied by rapid calls. In the cases in which the pair member involved in the chase went out of sight and the calling stopped, we assumed the encounter was over after 30 seconds.

In total, we recorded 1517 vocal events. Among these 244 (16%) were missing the distance between the pair members, as we couldn’t localize one pair member. The missing data were mostly for solo songs (n = 228). This made up 32% of the solo songs in the dataset and were mostly male solo songs (n = 193). In any case, our results were robust under various assumptions about the missing data (See supplementary materials for robustness analyses).

### Statistical analysis

All statistical analyses were done in R version 4.4.2 (R Core Team, 2024). We first described the dataset and calculated temporal statistics for duets. To quantify overlap, we calculated the gap between the end of an initiation song and the start of an answer song or call bout. As we subtracted the start of the answer from the end of initiation, negative values indicated overlap and positive values indicated alternation. For the answering precision, we calculated variability in the reaction time (i.e., how much time it took for a answerer to start a reply from the initiation of the partner’s song) as the coefficient of variation. We then built hierarchical generalized additive models using the *gam* function from the package *mgcv* to test our hypotheses (Wood, 2017). For all models, we checked concurvity and the suitability of the number of basis functions using the *gam.check* and *concurvity* functions from the *mgcv* package. We inspected residuals using the *DHARMa* package (Hartig, 2017). We used the package *ggeffects* to compute and plot model estimates and perform pairwise comparisons (Lüdecke, 2018). For the parameter estimates, non-focal parameters were averaged over the factor levels or set to their mean value with the *margin* argument in *predict_response* function set to *“marginalmeans”*. P-values for the pairwise comparisons were Tukey adjusted.

We first modeled the probability of answering (0 = solo song, 1 = song or call duet) with a binomial distribution. The model included the distance between the pair members, presence of a conspecific song before the vocal event, initiator sex and two-way interactions of initiator with distance and presence of a conspecific song before the vocal event as parametric terms. We modeled the effect of time relative to aggressive encounter along with its interaction with the initiator by modeling separate smooths with thin plane regression spline for each level of the initiator variable (i.e. male and female). The model also included pair ID as the random intercept. With the missing data excluded, this model had 1273 observations. To estimate non-distance-related parameters with a complete dataset, we ran the same model without the distance parameter.

Following the finding that answering probability didn’t depend on time relative to an aggressive encounter (see Results), we also analyzed whether initiation rates depended on the time relative to an aggressive encounter, as increased duet rates without increased answering rates were previously suggested as evidence for joint resource defense function (Krause et al. 2026). For this analysis, we summed the number of initiations (unanswered or answered) for each sex over each observation minute to obtain initiations per minute. We calculated time relative to an aggressive encounter using the center point of each block, and coded it as zero if the block contained an aggressive encounter. We then modeled the relationship between these variables for each sex using a negative binomial distribution to account for overdispersion in the data.

The last binomial model had the probability of answering with a song among the duets (0 = call duet, 1 = song duet) as the response variable, and included distance between the pair members and the presence of a conspecific song before the vocal event as parametric terms. The effect of time relative to an aggressive encounter was included as a thin plane regression spline. The model also included pair ID as the random intercept. This model only included male-initiated duets because female-initiated call duets were rare (see Results). With female-initiated song duets and missing data excluded, this model had 691 observations.

## Results

### Description of the dataset and duet structure

We recorded 1517 vocal events comprising 729 solo songs (i.e., unanswered initiations), 365 duets with song answers, and 423 duets with call answers. Female-initiated duets were uncommon: females initiated only 3% of call duets (n = 13) and 19% of song duets (n = 69). Answering rate to female initiations was 29%, whereas it was 57% to male initiations. Initiation and answering rates also varied among the pairs (Figure 2A). Overall, 64% of female songs were produced as part of a duet, compared with 41% of male songs. Most duet answers overlapped with the initiating partner’s song (Figure 2B). Among male-initiated duets, 79% of female call answers (n = 327) and 66% of female song answers (n = 196) overlapped with the male’s song.

**Figure 2.**
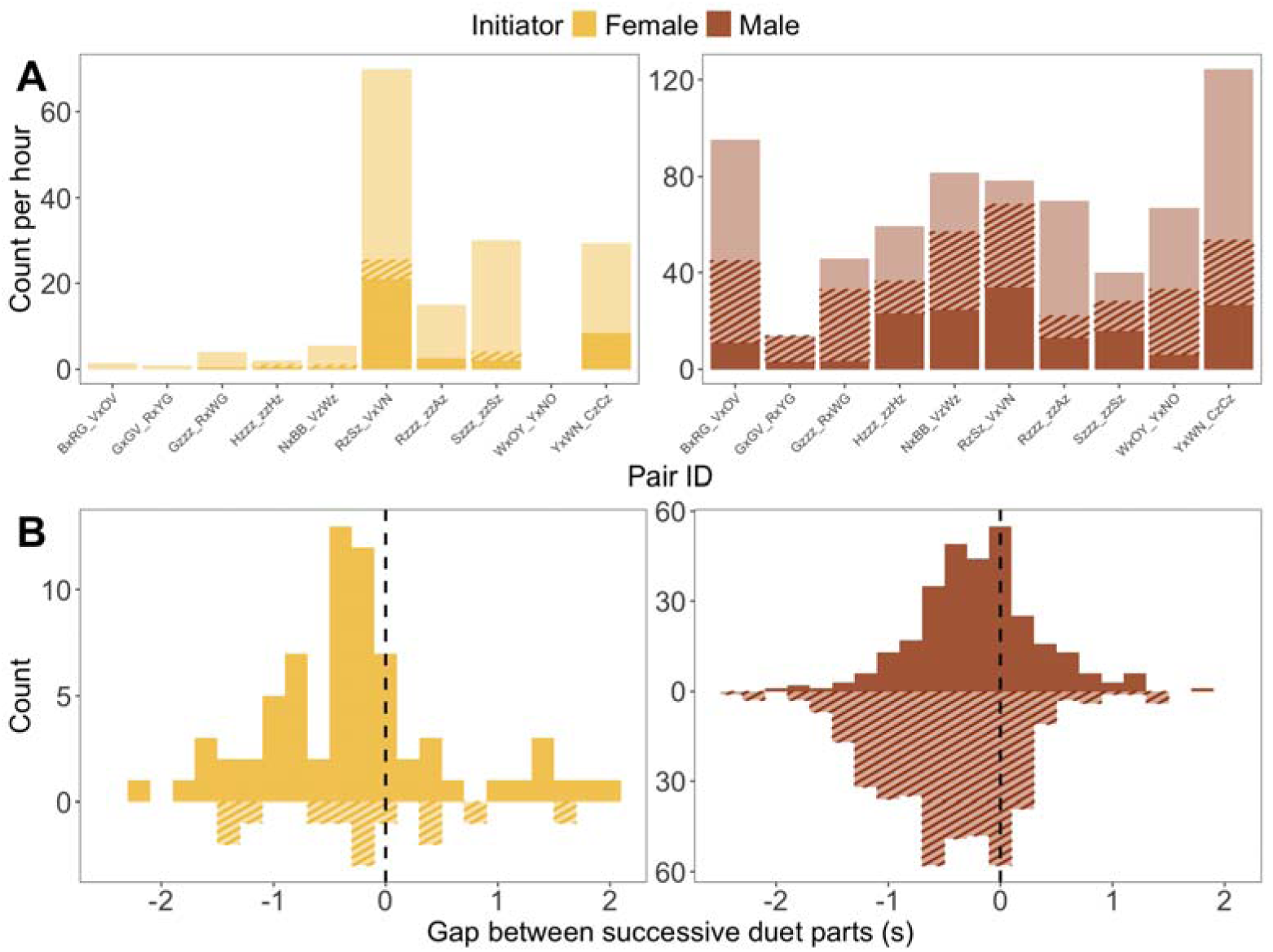
Hourly duet rates and temporal organization of duets A) Duet initiation and answering rates by sex and within pairs. Bars show the number of duet initiation attempts, the solid portions show the amount that the mate answered with a song and the patterned portions show the amount that the mate answered with a call bout. Note the different y-axis scales for females and males. B) Gap between the offset of an initiator’s song and onset of an answer; positive values indicate alternation while negatives show overlap. Similar to the plot above, the solid portions show song answers and the patterned portions show call answers.

Likewise, among female-initiated duets, 69% of male call answers (n = 9) and 70% of male song answers (n = 53) overlapped with the female’s song. Female answers were more temporally precise than male answers. The variability in female answering latency was 0.45 for call answers and 0.34 for song answers, compared with 0.75 and 0.57, respectively, for male answers.

### Context of answering

The probability of answering to a partner’s initiation depended on the interaction between the distance between the pair members and the initiator’s sex (see Table 1 and Figure 3A). Females were more likely to answer to male initiations when pair members were close together, whereas male answers to female initiations were virtually unaffected by distance. For male-initiated duets, the estimated probability of a female answering declined from 90% (95% CI = 0.86-0.92) at 1 m to 48% (95% CI = 0.41-0.55) at 22 m. In contrast, male answering probability to female initiations changed only slightly with distance, decreasing from 35% (95% CI = 0.24-0.48) at 1 m to 25% (95% CI = 0.15-0.38) at 22 m. This interaction effect remained robust even when we analyzed the data by assigning hypothetical values to missing data at both ends of the naturalistic distance range (Supplementary Figure 1, Supplementary Tables 1 and 2).

**Figure 3.**
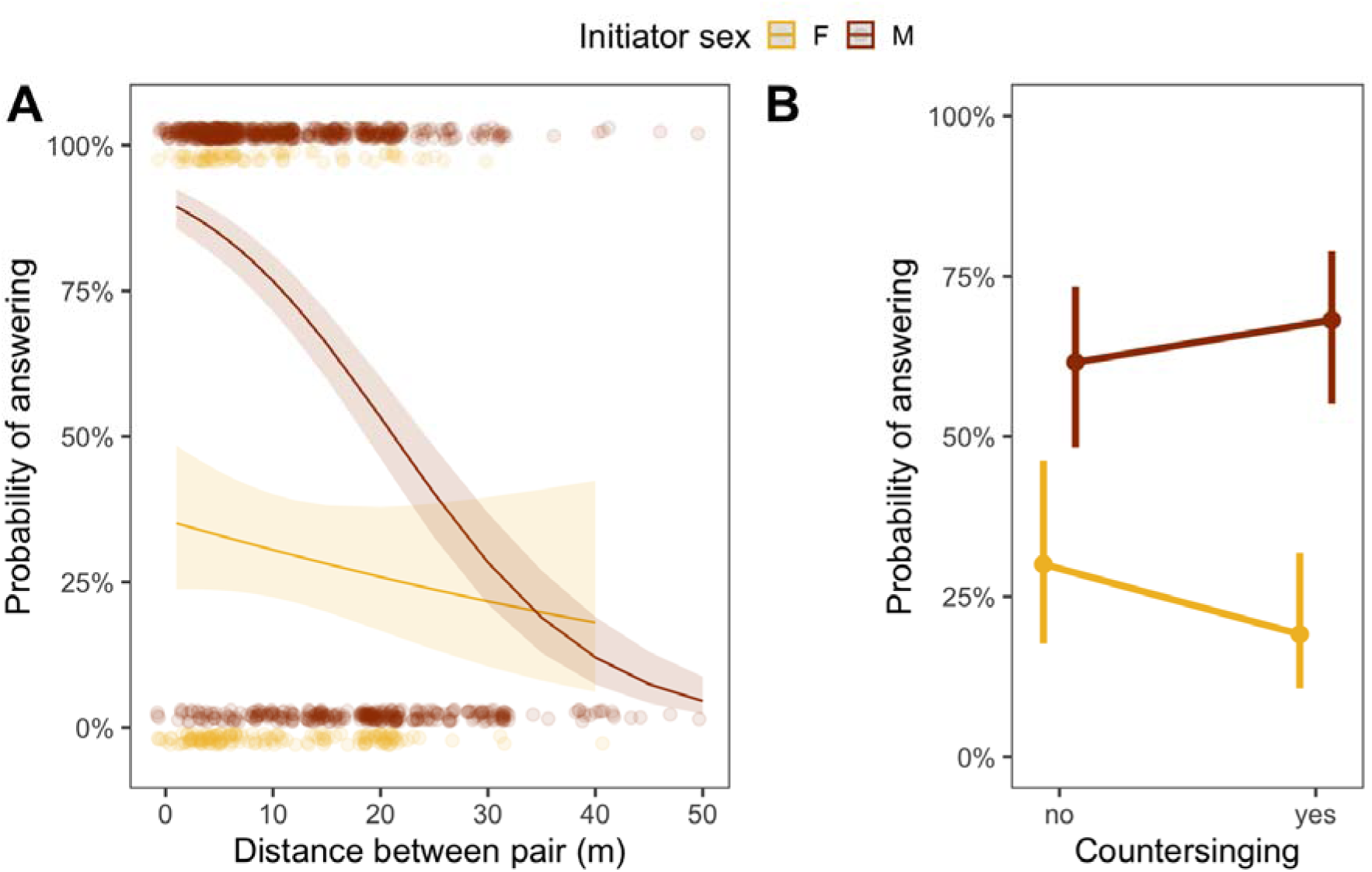
Effect of the distance between the pair members and countersinging depended on the sex for duet answering. A) Probability of answering with a song or call for each sex in relation to distance between pair members. Circles represent individual vocal events, and lines show the estimates with 95% confidence intervals shaded. B) Probability of answering with a song or call for each sex in relation to countersinging (i.e., whether a conspecific sang before the event). Circles with whiskers represent estimates with 95% confidence intervals.

**Table 1.**
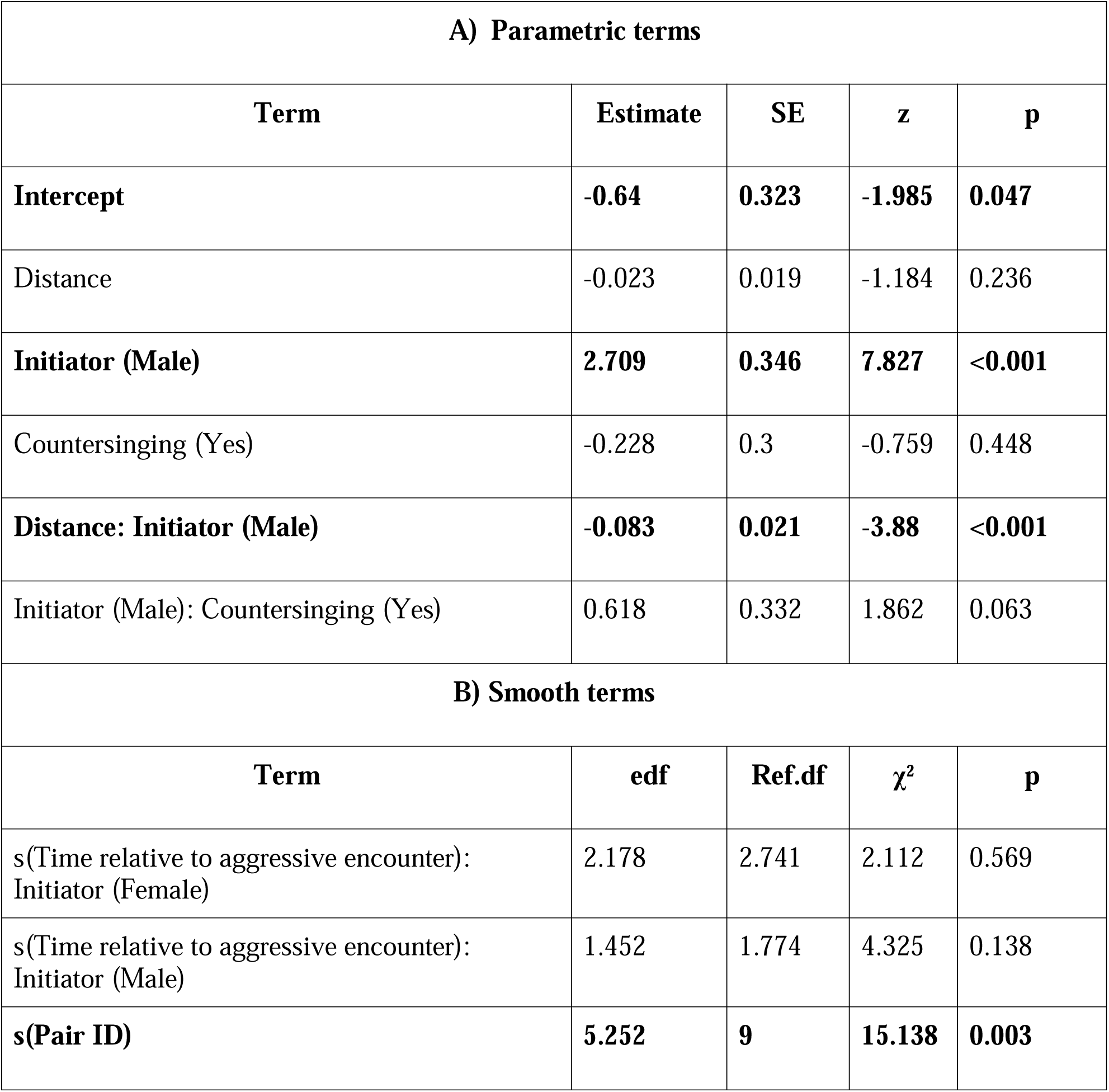
Results of the hierarchical generalized additive model for the probability of answering a duet initiation (either with a song or a call). Observations with missing distance values omitted.

**Table 2.** Results of the hierarchical generalized additive model for the probability of answering a duet initiation (either with a song or a call). To include all observations, distance is dropped from the previous model.

| **B) Parametric terms** |
| --- |

| **Term** | **Estimate** | **SE** | **z** | **p** |
| --- | --- | --- | --- | --- |
| **Intercept** | **-0.989** | **0.342** | **-2.894** | **0.004** |
| **Initiator (Male)** | **1.388** | **0.242** | **5.746** | **<0.001** |
| **Countersinging (Yes)** | **-0.597** | **0.297** | **-2.012** | **0.044** |
| **Initiator (Male): Countersinging (Yes)** | **0.887** | **0.316** | **2.803** | **0.005** |
| **B) Smooth terms** |  |  |  |  |
| **Term** | **edf** | **Ref.df** | **$\chi^2$** | **p** |
| s(Time relative to aggressive encounter): Initiator (Female) | 2.045 | 2.579 | 1.721 | 0.641 |
| s(Time relative to aggressive encounter): Initiator (Male) | 3.557 | 4.445 | 7.444 | 0.15 |
| **s(Pair ID)** | **8.116** | **9** | **123.044** | **<0.001** |

Although there was a non-significant trend in the first model with missing data (Table 1), the probability of answering also depended on the interaction between the presence of a conspecific song before the vocal event and the initiator’s sex when the complete dataset was analyzed (see Table 1). Apart from the slight change in p-value, estimates were comparable across models (See Supplementary Figure 2). For male initiations, the probability of a female answering was slightly higher when a neighbor conspecific sang before (68%, 95% CI = 0.55-0.79) than when there were no songs before (62%, 95% CI = 0.48-0.73; contrast = 0.07, 95% CI = 0.004-0.13, *P* = 0.04). There was a negative non-significant trend in males answering to a female-initiated song, depending on the presence of a neighbor song prior to the vocal event (contrast = −0.11, *P* = 0.057).

Answering probability didn’t depend on the time relative to an aggressive encounter for the initiations of either sex (see Table 1). Inspecting the initiation rates to test if the duet output changes with time relative to an aggressive encounter also revealed no relationship for either sex (see Supplementary Figure 3 and Supplementary Tables 3-4).

### Context of the song answering

Among male-initiated duets, the probability that a female would answer with a song depended on the distance between the pair members (see Table 3, Figure 4A). Females were equally likely to answer with a song or a call at close distances (predicted probability at 1m = 0.53, 95% CI = 0.37-0.68); however, when they were farther apart, they were less likely to answer with a song (predicted probability at 20m = 0.30, 95% CI = 0.18-0.45). Although the fitted smooth for the effect of time relative to an aggressive encounter was significantly different from a flat response, this effect is unlikely to be behaviorally relevant, as there was no change in probability within a five-minute window relative to an aggressive encounter (see Table 3 and Figure 4B). Females also showed a weak, non-significant tendency to answer with a song rather than a call when a neighboring conspecific had sung beforehand (Table 3, Figure 4C; contrast = 0.09).

**Figure 4.**
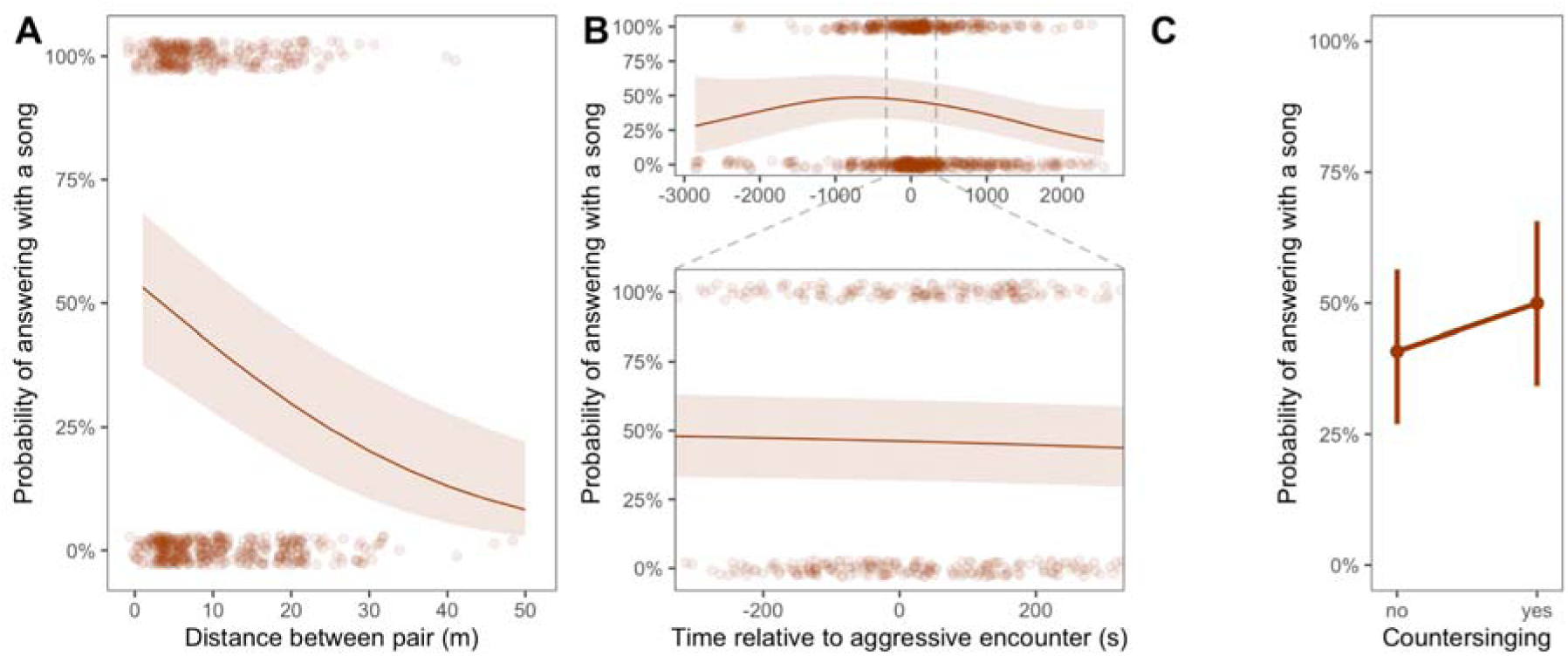
In male-initiated duets, the probability of a female answering with a song depended on the distance between pair members and time relative to an aggressive encounter. Furthermore, there was a non-significant trend for females being slightly more likely to answer with a song in countersinging with neighbors. **A)** Probability of answering with a song in duets was higher when the pair was closer to each other. **B)** Although the effect of time relative to the aggressive encounter was statistically significant in distinguishing song vs. call answers, this effect is unlikely to be ecologically relevant with respect to our hypotheses, as the probability curve was flat within the five minutes of an encounter. In both figures, circles represent individual vocal events, and lines show the estimates with 95% confidence intervals shaded. **C)** Probability of answering with a song among duets in relation to countersinging (i.e., whether a conspecific sang before the event). Circles with whiskers represent estimates with 95% confidence intervals.

**Table 3.** Results of the hierarchical generalized additive model for the probability of answering with a song among duets.

| **A) Parametric terms** |  |  |  |  |
| --- | --- | --- | --- | --- |
| **Term** | **Estimate** | **SE** | **z** | **p** |
| Intercept | -0.117 | 0.333 | -0.35 | 0.726 |
| **distance** | **-0.052** | **0.012** | **-4.205** | **<0.001** |
| conspecific_song_beforeeyes | 0.366 | 0.198 | 1.848 | 0.065 |
| **B) Smooth terms** |  |  |  |  |
| **Term** | **edf** | **Ref.df** | **$\chi^2$** | **p** |
| **s(Time relative to aggressive encounter)** | **2.597** | **3.281** | **11.31** | **0.0124** |
| **s(Pair ID)** | **7.850** | **9.000** | **60.85** | **<0.001** |

## Discussion

In this study, we took an observational approach to understand the sex-specific functions of answering in duets and whether song and call answers have different functions. We found that male initiations received higher answer rates than female initiations. Although both sexes answered to their partners’ initiations with songs, males rarely answered female initiations with *chip* calls. Moreover, females answered more precisely than males with higher temporal coordination in their replies, especially when answering with songs rather than calls. The influence of context on answering behavior also differed between the sexes. Among the three contexts we investigated, distance to the mate had the strongest effect on female answering behavior, followed by a slight effect of countersinging with neighbors, whereas aggressive encounters had no biologically meaningful effect. In contrast, none of the investigated contexts substantially influenced male answering behavior.

An abundance of duetting species in tropical habitats with dense vegetation led to the hypothesis that duet initiation and answers enable contact maintenance between the pair members (Thorpe 1963). Under this hypothesis, we expected pair members to be further apart from each other (i.e., without visual contact). In contrast to this hypothesis, while male answering behavior did not vary meaningfully with distance to partner, female answering was more likely when they were already closer to their mate. Moreover, at close distance, females were equally likely to answer with a song or a call; however, the predicted probability of singing dropped with increased distance to the mate. Therefore, our results do not support the contact maintenance hypothesis for duetting. Some studies found partial support for the hypothesis, showing that pair members were more likely to approach each other after duetting (Logue 2007; Mennill and Vehrencamp 2008), but we were unable to test this possibility in our dataset. Nevertheless, the same studies found that pair members were already close to each other when they answered their mate, suggesting contact maintenance is not likely a primary function. Even if Galápagos Yellow Warblers approached one another after duetting, this behaviour would not explain our main finding that females answered most frequently when already close to their mates, rather than when separated, as predicted by the contact maintenance hypothesis. Overall, our findings provide little support for contact maintenance as the primary function of duetting in this species.

We also tested the joint resource defense hypothesis, which is the best-supported hypothesis for duetting to date (Dahlin and Benedict 2014). According to this hypothesis, duets are joint signals to extrapair receivers that advertise and/or defend territory ownership (Hall 2004). For the latter, we tested whether answering the mate was associated with agonistic encounters inside the territory. We predicted that answering would be more likely during, or just before or after, an agonistic interaction to defend the territory against intruders. Our results showed no strong evidence of an association. Among female answers, the probability of singing varied with respect to the time of an aggressive encounter, peaking at about 15 minutes before an actual aggressive interaction. However, there was no such difference within a more immediate time scale of 5 minutes before or after (Figure 4B). We therefore interpret these findings as providing little support for the joint resource defense hypothesis, as a joint defense signal would be expected to occur most frequently during, or immediately after, an aggressive encounter.

Several other studies also reported that answering rates do not vary with respect to agonistic interactions (Fedy and Stutchbury 2005; Diniz et al. 2018; Krause et al. 2026), however, there seems to be a lack of consensus on interpretation. Krause and colleagues (2026) argue that if duet rates increase during an agonistic interaction, even as a byproduct of increased initiation rates without increased answer rates, this should be taken as evidence for the joint resource-defense function of duetting. In our view, the evidence for the joint defense function would be rather weak, because any function requiring a stable answer rate, such as signalling attentiveness to the partner, would produce the same pattern. We therefore suggest that only an increase in answer rate provides convincing evidence for the joint resource defense function. Nevertheless, receivers may attend to changes in duet rates rather than answer rates. Disentangling these would be a fruitful avenue for future research. In any case, our analyses suggest that initiation rates, and therefore duet rates didn’t change with respect to their relative time to an agonistic encounter. Moreover, our previous experiments showed that neither sex perceived duet playbacks at a 100% answer rate as more threatening than solo songs (Yelimlieş et al., 2026), making Galápagos Yellow Warblers unsuitable for such research. Overall, we believe that there is sufficient evidence suggesting that duets are not used in agonistic interactions during territorial defense in this species.

Alternatively, answers from either partner to an initiation may signal joint ownership of the territory, which could be useful during vocal exchanges with neighboring pairs. We tested whether answering to a partner changed depending on whether the partner was countersinging with a neighboring conspecific. We found some support for this version of the hypothesis, as female answer rates increased slightly when a neighbor sang before the mate. Moreover, there was a trend indicating female answers in duets were more likely to be songs in a countersinging context, yet this effect was not statistically significant. Using duets in disputes with neighbors has been observed in several other species (Harcus 1977; Benedict 2010; Odom et al. 2017). Nevertheless, the predicted increase in answers was only 7%, and we cannot rule out the possibility that this is not a biologically meaningful change. A promising way to test the significance of duets in signaling joint ownership is to conduct mate- or pair-removal experiments (Krebs et al. 1978; Levin 1996c). If duets function as signals of joint territory ownership, they should deter intruders more effectively than solo songs.

Our findings didn’t reveal a significant relationship between the male answering behavior and neighbour singing, but the trend was in the opposite direction, with males being less likely to answer when the female song was preceded by a neighbour song. We believe this finding is consistent with several other findings from our study. We showed that males initiated the majority of duets, and male songs were more likely to be answered by the mate than female songs. Additionally, we found that male answers were less temporally coordinated than female answers. While the variation in female reaction times was 34% for song and 45% for call answers, the variation in male song reaction times was 57%. To put these numbers in context, while female Galápagos yellow warblers’ song coordination is far more variable than some antiphonal duetters that have below 10% variation (Levin 1996c; Hall and Magrath 2007), they are more coordinated than some species like Superb Fairywrens (*Malurus cyaneus*; 54% variation that seemingly shows just above random patterns in answering Taylor et al., 2019) and closer to some other species like Eastern Whipbirds (*Psophodes olivaceu*s; Rogers, 2005) with 25% variation. Taken together, the high level of variation in male song answering times and the low number of female-initiated duets suggest that male answering behavior in this species may not be a biologically meaningful phenomenon.

The main goal of our study was to understand why song answering in females evolved by uncovering its adaptive function. Among the contexts we investigated, distance to the mate seemed to be the main driver of singing rather than calling in female answers. Contrary to what the contact maintenance hypothesis would predict, females were less likely to sing as an answer to male initiations when they were further apart from their mate.On the other hand, females singing when close to their mates is consistent with one prediction of the joint resource defense hypothesis. By being close to each other while signaling, the pair poses a greater threat to potential intruders than distantly positioned single individuals (Hultsch and Todt 1984). However, because we found only a non-significant trend towards increased song answers during countersinging with neighbours, and no meaningful association with time relative to aggressive encounter, support for this explanation remains limited.

An alternative, non-mutually exclusive explanation for the evolution of female song answers in duets is that they signal pair quality or commitment (Hall 2004). By answering quickly and consistently, females may signal attentiveness and commitment to the pair bond to the mate and extrapair receivers. This idea is supported by the fact that female song answers had the smallest variation in reaction times. Furthermore, shorter reaction times would require being closer to the mate, which is consistent with females being close to their mates when they sing to answer as opposed to calling. Another remaining question then is why females answer with songs rather than calls to maintain the pair bond. One possible proposed function is that mates learn and use each other’s duet contributions, and learning investment makes abandoning the pair bond costly (Thorpe and North 1965); however, this is unlikely to be relevant for Galápagos Yellow Warblers, as duet contributions are sex-specific (Figure 1A). Instead, we speculate that female song may have higher potential for signaling individual identity than chip calls, although we couldn’t test this hypothesis here due to insufficient good-quality recordings. If that is the case, females could signal their commitment to the pair bond to extrapair receivers. Lastly, we argue that the pair bond commitment hypothesis would explain the sexual asymmetry in duet formation. Females could show higher effort by answering at a higher rate and more precisely than males because they could be at a higher risk of losing their mate. Indeed, we have observed facultative polygyny, but not polygamy, in our population.

In conclusion, the present study adds to the growing literature on the form and functions of avian duetting. As in many other species, Galápagos Yellow Warbler duets were mostly female-answer-driven and produced at short distances between the pair. We found limited support linking female answers to the joint resource defense hypothesis. On the other hand, we argue that duetting and, more specifically, the evolution of song answers could be potentially explained by a need to signal commitment to the pair bond. Although this idea requires testing, it is consistent with the selection pressures that would arise from transitioning from a migratory to a sedentary life history, which happened during the split of Galápagos Yellow Warblers (subspecies of Mangrove Yellow Warbler) and Northern Yellow Warbler.

## Acknowledgements

We are grateful to the Galápagos National Park Directorate for permission to conduct this study (permit numbers PC-87-23 and PC-05-25) and the Charles Darwin Research Station for logistical support. We are especially grateful to the Galápagos National Park Directorate on Floreana, which has been a cornerstone of this long-term research project. In particular, we thank Eddie Rocero, Director of the Galápagos National Park on Floreana during this study, and Francisco Moreno for their longstanding support and collaboration. We also thank the Floreana community, especially Claudio Cruz, Walter Cruz, Erica Wittmer, and Ingrid Wittmer, for their generous support throughout many years of fieldwork. We thank Jefferson García Loor for logistical support, Lauren K. Common for organising the long-term project data, and both Lauren K. Common and Jefferson García Loor for leading the field team during bird banding. Finally, we acknowledge the dedication of the many members of the Floreana Ornithology Team, whose collaborative work across projects has made this long-term research programme possible. This publication is contribution number XXXX (to be filled upon acceptance) of the Charles Darwin Foundation for the Galápagos Islands.

## Funding

This study is funded by the British Ornithologists’ Union Small Ornithological Research Grant awarded to AY, and by the Austrian Science Fund (project numbers W1262-B29 and P 36342-B) with awards to SK.

## Conflicts of interest

The authors declare no conflict of interest.

## Data availability

Analyses reported in this article can be reproduced using the data provided at https://osf.io/n9d4g/overview?view_only=9ff74bdaf0fc4756962700290556ccdc

## Notes

https://osf.io/n9d4g

### Competing Interest Statement

The authors have declared no competing interest.

