## Supplementary Materials for "Sex-specific functions of duetting in the Galápagos Yellow Warbler (Setophaga petechia aureola)"

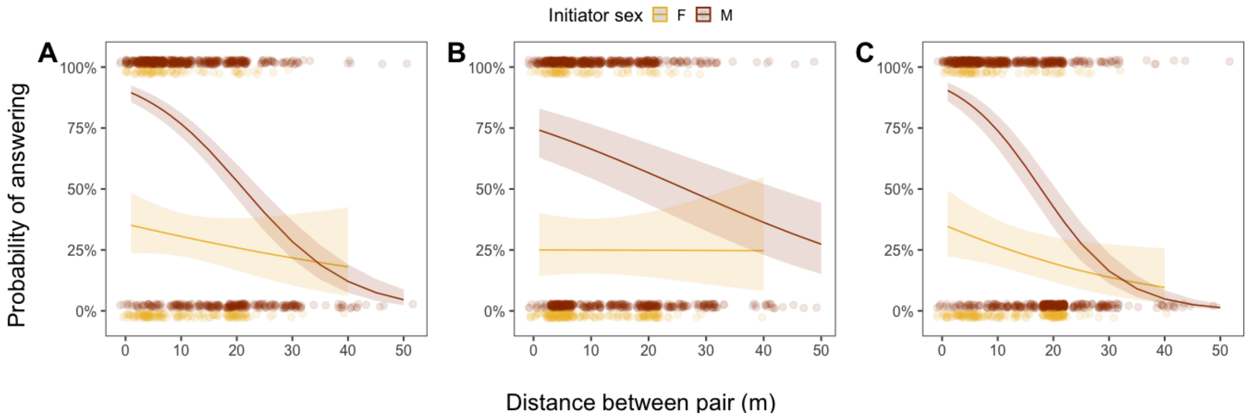


**Supplementary Figure 1.** The interaction effect between sex and distance on the probability of answering a duet initiation doesn't depend on the excluded observations due to missing distance values. **A)** Model predictions with missing values excluded, same figure as in the original text. **B)** Model predictions with all the missing distance values set to 5 m (1st quartile of the data). **C)** Model predictions with all the missing distance values set to 20 m (3rd quartile of the data). Note that missing values arise from the failure to locate one pair member as the other sings. Therefore, we argue that the scenario simulated in B is less likely than C.

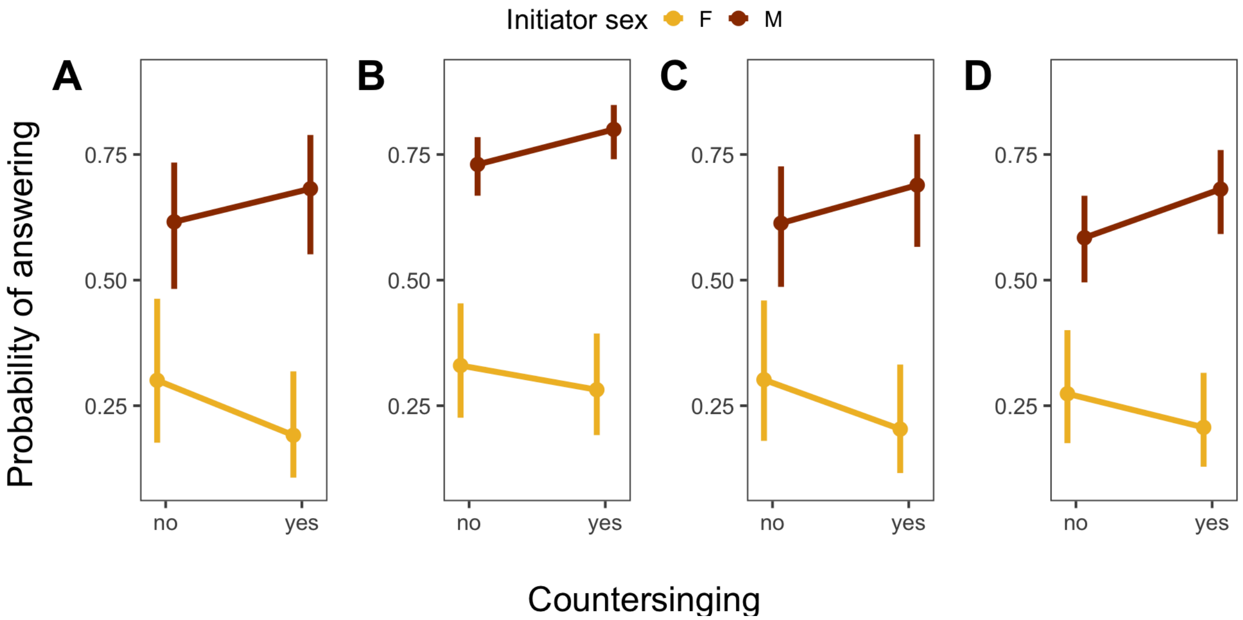


**Supplementary Figure 2.** Including observations with missing distance values yields similar estimates of the probability of duet answering for each sex, depending on countersinging; however, the statistical significance of the interaction term depends on their inclusion (see Tables 1 & 2 in the main text and the Supplementary Tables 1 & 2). **A)** Model predictions from the model without the distance parameter that includes all observations, same figure as in the original text. **B)** Model predictions from the model with missing values excluded had a non-significant interaction term (See Table 1). **C)** Model predictions with all the missing distance values set to 5 m. **D)** Model predictions with all the missing distance values set to 20 m.

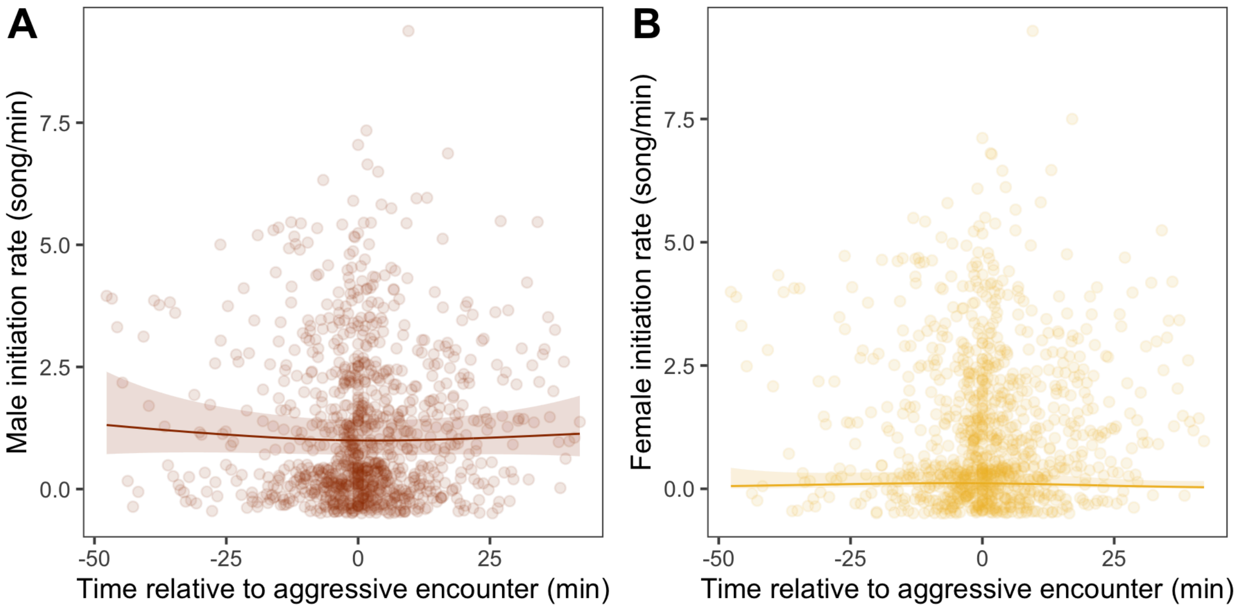


**Supplementary Figure 3.** Initiation rates did not depend on the time relative to aggressive encounter for either sex. A) Predicted number of male and B) female duet initiations per minute depending on the number of neighbour songs during that minute. Circles represent each minute, and lines show the estimates with 95% confidence intervals shaded.

**Supplementary Table 1.** Results of the hierarchical generalized additive model for the probability of answering, with missing distance values set to 5 m.

| **A) Parametric terms** | | | | |
|---|---|---|---|---|
| **Term** | **Estimate** | **SE** | **z** | **p** |
| **Intercept** | **-1.021** | **0.386** | **-2.647** | **0.008** |
| Distance | -0.00045 | 0.02 | -0.023 | 0.982 |
| **Initiator (Male)** | **1.869** | **0.325** | **5.75** | **<0.001** |
| Countersinging (Yes) | -0.526 | 0.297 | -1.771 | 0.077 |
| **Distance: Initiator (Male)** | **-0.041** | **0.021** | **-1.966** | **0.049** |
| **Initiator (Male): Countersinging (Yes)** | **0.861** | **0.317** | **2.711** | **0.007** |
| **B) Smooth terms** | | | | |
| **Term** | **edf** | **Ref.df** | **$\chi^2$** | **p** |
| s(Time relative to aggressive encounter): Initiator (Female) | 2.321 | 2.919 | 2.216 | 0.559 |
| s(Time relative to aggressive encounter): Initiator (Male) | 3.847 | 4.787 | 8.713 | 0.12 |
| **s(Pair ID)** | **7.984** | **9** | **108.819** | **<0.001** |

**Supplementary Table 2.** Results of the hierarchical generalized additive model for the probability of answering, with missing distance values set to 20 m.

| **A) Parametric terms** | | | | |
|---|---|---|---|---|
| **Term** | **Estimate** | **SE** | **z** | **p** |
| **Intercept** | **-0.591** | **0.345** | **-1.712** | **0.087** |
| **Distance** | **-0.041** | **0.019** | **-2.185** | **0.029** |
| **Initiator (Male)** | **2.759** | **0.349** | **7.898** | **<0.001** |
| Countersinging (Yes) | -0.369 | 0.295 | -1.25 | 0.211 |
| **Distance: Initiator (Male)** | **-0.092** | **0.021** | **-4.42** | **<0.001** |
| **Initiator (Male): Countersinging (Yes)** | **0.787** | **0.321** | **2.454** | **0.014** |
| **B) Smooth terms** | | | | |
| **Term** | **edf** | **Ref.df** | **$\chi^2$** | **p** |
| s(Time relative to aggressive encounter): Initiator (Female) | 2.231 | 2.804 | 2.324 | 0.528 |
| s(Time relative to aggressive encounter): Initiator (Male) | 1.001 | 1.003 | 2.745 | 0.098 |
| **s(Pair ID)** | **6.917** | **9** | **44.327** | **<0.001** |

**Supplementary Table 3.** Results of the hierarchical generalized additive model for the male duet initiation rates per minute.

| **B) Parametric terms** | | | | |
|---|---|---|---|---|
| **Term** | **Estimate** | **SE** | **z** | **p** |
| **Intercept** | 0.0156 | 0.1761 | 0.089 | 0.929 |
| **B) Smooth terms** | | | | |
| **Term** | **edf** | **Ref.df** | **$\chi^2$** | **p** |
| s(Time relative to aggressive encounter) | 1.787 | 2.263 | 1.714 | 0.487 |
| **s(Pair ID)** | **8.36** | **9** | **86.43** | **<0.001** |

**Supplementary Table 4.** Results of the hierarchical generalized additive model for the female duet initiation rates per minute.

| **C) Parametric terms** | | | | |
|---|---|---|---|---|
| **Term** | **Estimate** | **SE** | **z** | **p** |
| **Intercept** | **-2.297** | **0.559** | **-4.11** | **<0.001** |
| **B) Smooth terms** | | | | |
| **Term** | **edf** | **Ref.df** | **$\chi^2$** | **p** |
| s(Time relative to aggressive encounter) | 2.33 | 2.98 | 6.21 | 0.097 |
| **s(Pair ID)** | **8.25** | **9** | **193.49** | **<0.001** |
